# Robust Reference Powered Association Test of genome-wide association studies

**DOI:** 10.1101/316703

**Authors:** Yi Wang, Yi Li, Meng Hao, Xiaoyu Liu, Menghan Zhang, Momiao Xiong, Jiucun Wang, Yin Yao Shugart, Li Jin

## Abstract

Genome-wide association studies (GWAS) have identified abundant genetic susceptibility loci, although they are far less from meeting the previous expectations due to low statistical power and false positive results. Effective statistical methods are required to further improve the analyses of massive GWAS data. Here we presented a new statistic (Robust Reference Powered Association Test, http://drwang.top/gwas.html) to use large public database as reference to reduce concern of potential population stratification. To evaluate the performance of this statistic for various situations, we simulated multiple sets of sample size and frequencies to compute statistical power. Furthermore, we applied our method to several real datasets (psoriasis genome-wide association datasets and schizophrenia genome-wide association dataset) to evaluate the performance. Careful analyses indicated that our newly developed statistic outperformed several previously developed GWAS applications. Importantly, this statistic is more robust than naive merging method in the presence of small control-reference differentiation, therefore likely to detect more association signals.

## INTRODUCTION

Genome-wide association studies (GWAS) have been widely applied with the goals to detect genetic variants which contribute to complex traits in the past decade (1). In general, allele frequencies of genetic variants are compared between cases that are supposed to have a high prevalence of susceptibility alleles and controls that are considered to have a lower prevalence of such alleles. And genomic loci correlated with various traits had been detected using the efficient approaches (2).

Although GWASs have led to abundant significant findings (3-7), a few practical difficulties hinder the discovery of more rare or low-frequency genetic variants. For example, limited sample sizes make it difficult to achieve high statistical power which shows the probability of identifying the latent genetic variants (2,8). Besides, difference in genetic background, also known as population stratification, between cases and controls could inflate type I error rate, thereby, leading to increasing level of false positive findings (9).

Availability of large public datasets contain large amount of useful genetic information. Utilizing large public datasets as reference may increase the sample size and ease the low power brought by insufficient sample size. However, population stratification and genotyping platform differences (10) between public datasets and cases may lead to inflated type I error rate. Our study aim is to appropriately utilize the public datasets to conservatively ameliorate the situation of small samples and low statistical power. In this manuscript, we describe a novel method to properly use public datasets as reference. In particular, we introduce a large public population as reference which has similar genetic background with control group. The null hypothesis was that allele frequency of SNPs among case (ca), control (co) and reference (re) was equal, and that the control and reference might compose a new large reference (co+re).

Specifically, we constructed a new test statistic *T*=*ln*(*p*(*ca*–(*co*+*re*)))–*ln*(*p*(*co*–*re*)) in which *p*(*ca*–(*co*+*re*)) is p-value of 2×2 Fisher exact test (11-13) of the SNP between case and control+reference, and *P* (*co*–*re*) is p-value of 2×2 Fisher exact test (11-13) of the SNP between control and reference. The T statistic takes differences between control and reference into account, which is more robust than *p*(*ca*–(*co*+*re*)) or *p*(*ca*–*re*).

In this manuscript, we present a method to use public datasets as reference in the association analysis. The results of simulation and real data application showed that our new method could increase statistical power. And the online tool (Robust Reference Powered Association Test, http://drwang.top/gwas.html) has been made available.

## MATERIAL AND METHODS

### Framework of Robust Reference Powered Association Test

Suppose we have three populations: case, control and public data pools, intuitively we want to merge the control and reference population to form a large control pool to gain more power on allele-disease association test. However, we are concerned about the potential population differentiation and genotyping platform difference between control and the reference (10).

A simplistic way to alleviate such concern is to perform a control (co) vs. reference (re) Fisher exact test (11). If a p-value is not significant and the control sample size is not too small, then this concern is resolved. However, choosing the significance level is arbitrary and the decision is subjective. We need an objective version of such procedure to correct the effect of co-re (control vs. reference) difference.

Denote the p-value of 2×2 Fisher exact test (11-13) of the SNP between case and control+reference as *P*(*ca*–(*co*+*re*)). Denote the p-value of 2×2 Fisher exact test of the SNP between control and reference as *p*(*co*–*re*). We define a test statistic: *T*=*ln*(*p*(*ca*–(*co*+*re*)))–*ln*(*p*(*co*–*re*))(Figure 1). T will be smaller if the difference between case and control + reference is larger, while T will be larger if the difference between control and reference is larger. Therefore, T is a more robust statistic than *p*(*ca*–(*co*+*re*))as it takes the control-reference difference into account by penalizing *p*(*co*–*re*).

**Figure 1.**
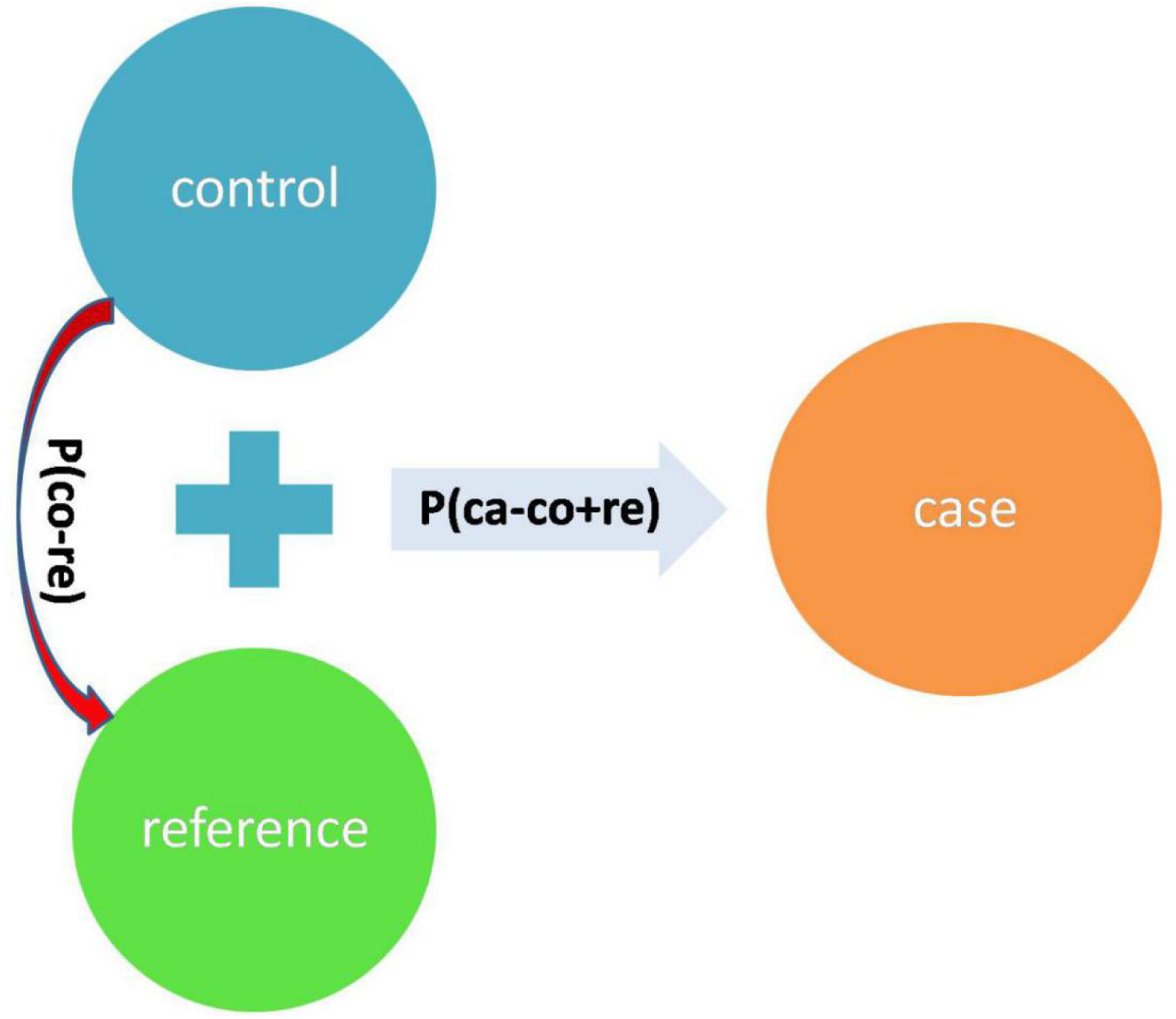
The summarized design of the test statistic T.

Our null hypothesis is that the allele frequency of the SNP is equal among cases, controls and references. Under the null hypothesis, both *p*(*ca*–(*co*+*re*)) and *p*(*co*–*re*))are independent and uniformly distributed on (0, 1]. Under the null hypothesis, –*ln(p*(*ca*–(*co*+*re*))) and –*ln(p*(*ca*–*co*))are exponentially distributed with parameter 1 (14). And T is the difference of two exponentially distributed variables, thus T is Laplace distributed as T~ Laplace (0, 1) (15). The one side p-value of T is as follow:

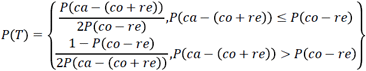

From the above formula one can easily see that the *p*(*co*–*re*)acts as a penalizer/normalizer against *p*(*ca*–(*co*+*re*)). Therefore, our new statistic T is more robust than the simple *p*(*ca*–(*co*+*re*))method in the presence of small control-reference differentiation. However, in the presence of strong population differentiation or genotyping platform difference, even our correction may not be effective, we therefore need to restrict *P* (*co*–*re*) to be not significant with a suggested threshold of 0.01.

### The psoriasis GWAS datasets

We obtained the psoriasis datasets (16, 17), as a part of the Collaborative Association Study of Psoriasis (CASP), from the Genetic Association Information Network (GAIN) database (dbGaP accession number: phs000019.v1. p1), a partnership of the Foundation for the National Institutes of Health. All genotypes were filtered by checking for data quality. A dermatologist diagnosed all psoriasis (MIM: 177900) cases. Each participant’s DNA was genotyped with the Perlegen 500K array. Both cases and controls agreed to sign the consent contract, and controls (≥18 years old) had no confounding factors relative to a known diagnosis of psoriasis.

### The schizophrenia GWAS dataset

The schizophrenia dataset (18) came from the GAIN dataset (dbGaP access number: phs000021.v1.p1), including 1,195 cases with schizophrenia (MIM: 181500) and 954 controls. The subjects were genotyped on AFFYMETRIX AFFY_6.0 platform. All subjects were at least 18 years old. The cases included 746 males (41.9 ± 10.8 years) and 449 females (43.0 ± 9.8 years); and the controls included 362 males (46.2 ± 13.7 years) and 592 females (45.0 ± 12.9 years). Affected subjects met lifetime DSM-IV criteria for schizophrenia (American Psychiatric Association 1994). Cases were excluded if they had worse than mild mental retardation, or if their psychotic illness was judged to be secondary to substance use or neurological disorders. Controls were excluded if they did not deny all of the following psychosis screening questions: treatment for or diagnosis of schizophrenia or schizoaffective disorder; treatment for or diagnosis of bipolar disorder or manic-depression; treatment for or diagnosis of psychotic symptoms such as auditory hallucinations or persecutory delusions.

## RESULTS

### Results from simulation study

To evaluate the performance of our model for various situations, we simulated six parameters to compute the desired statistical power. The parameters include case sample size, allele frequency in case, control sample size, allele frequency in control, reference sample size and allele frequency in reference. Six different parameters were set to several typical values to simulate real scenarios. The case and control size were set from small to large. Also, allele frequencies were set from rare to common. Additionally, we selected two different reference of 10,000 and 100,000 samples. In detail, we set case size (case = 100, 500, 1000, 3000), control size (control = 0.5*case, 1*case, 2*case, 5*case), reference size (reference = 10,000, 100,000). And allele frequencies were set in the reference (ref = 0.001, 0.01, 0.05, 0.15, 0.3), in control (frequencies =1*ref, 1.1*ref) and in case (frequencies =1*ref, 1.1*ref, 1.5*ref, 3*ref). Totally, there were 128,000 different combinations (see Additional files 1).

We have selected several representative cases to analyze the whole results of simulations (Table 1). First, supposed that the sample size of case, control and reference equaled 500, 500 and 10,000 respectively which were close to the real conditions. And allele frequencies in case, control and reference were initialized to be common (0.15). As indicated in Table 1, the power of T was a bit higher than power of ca-(co+re), while both were less than false positive rate (alpha=0.05, 0.01, 0.001). When there was small population genetic differentiation between control and reference, allele frequencies in case and control equaled 0.165 while allele frequency in reference was set as 0.15. In this case, the power of T was less than the power of ca-(co+re). Therefore, as mentioned in Method, our new test is more robust than the simple *p*(*ca*–(*co*+*re*))method in the presence of small control-reference differentiation.

**Table 1.** Simulation power in representative cases.

| Groups | N | Frequency | Alpha | ca-(re+co) | co-re | ca-co | T |
| --- | --- | --- | --- | --- | --- | --- | --- |
| case | 500 | 0.150 | 0.050 | 0.0472 | 0.0463 | 0.0431 | 0.0481 |
| control | 500 | 0.150 | 0.010 | 0.0092 | 0.0089 | 0.0079 | 0.0094 |
| ref | 10000 | 0.150 | 0.001 | 0.0008 | 0.0008 | 0.0008 | 0.0008 |
| case | 500 | 0.165 | 0.050 | 0.2290 | 0.2505 | 0.0426 | 0.1465 |
| control | 500 | 0.165 | 0.010 | 0.0895 | 0.0998 | 0.0080 | 0.0568 |
| ref | 10000 | 0.150 | 0.001 | 0.0193 | 0.0223 | 0.0008 | 0.0118 |
| case | 500 | 0.165 | 0.050 | 0.2142 | 0.4228 | 0.0478 | 0.0979 |
| control | 1000 | 0.165 | 0.010 | 0.0807 | 0.2098 | 0.0094 | 0.0359 |
| ref | 10000 | 0.150 | 0.001 | 0.0182 | 0.0655 | 0.0009 | 0.0078 |
| case | 500 | 0.165 | 0.050 | 0.1771 | 0.7398 | 0.0485 | 0.0323 |
| control | 2500 | 0.165 | 0.010 | 0.0622 | 0.5127 | 0.0097 | 0.0109 |
| ref | 10000 | 0.150 | 0.001 | 0.0124 | 0.2509 | 0.0009 | 0.0021 |
| case | 500 | 0.165 | 0.050 | 0.2496 | 0.0464 | 0.1367 | 0.2383 |
| control | 500 | 0.150 | 0.010 | 0.0992 | 0.0091 | 0.0429 | 0.0949 |
| ref | 10000 | 0.150 | 0.001 | 0.0228 | 0.0009 | 0.0075 | 0.0215 |
| case | 500 | 0.225 | 0.050 | 1.0000 | 0.0465 | 0.9891 | 0.9998 |
| control | 500 | 0.150 | 0.010 | 0.9997 | 0.0092 | 0.9524 | 0.9990 |
| ref | 10000 | 0.150 | 0.001 | 0.9965 | 0.0009 | 0.8300 | 0.9941 |
| case | 500 | 0.030 | 0.050 | 0.9953 | 0.0418 | 0.8799 | 0.9919 |
| control | 500 | 0.010 | 0.010 | 0.9808 | 0.0076 | 0.7038 | 0.9742 |
| ref | 10000 | 0.010 | 0.001 | 0.9259 | 0.0008 | 0.4246 | 0.9152 |

As shown in Table 1, false positive results might occur if there was small population genetic differentiation between control and reference while no differentiation between case and control. To evaluate the influence of control size, we simulated different control sizes with the aim to detect the false positive rate under different allele frequencies of case and control (see Additional files 2). We found that the false positive rate of P(T) would decrease when control size was augmented. Furthermore, P(T)’s power was less than *p*(*ca*–*co*)’s when the sample size of the controls was five times larger than that of the cases. The false positive results were kept as low as possible when the sample size of the controls was four times larger than that of the cases (see Additional files 2).

When the allele frequency in case was different from that in control, T’s power was always higher than power of ca-co. We could find that when there was a slight change of allele frequency in case, the power of T was much higher than power of ca-co, indicating that our method had high sensitivity for GWAS. When the allele frequency in case was much higher than control’s, the power approached to 1 with remarkable increase of T’s power. When the allele frequency was rare, we could also draw the same conclusion that our method could keep false positive rate low and drastically increase the statistical power.

### Results from the psoriasis GWAS datasets

We applied our newly developed method to two psoriasis genome-wide association (GWAS) (16,17) datasets to evaluate the performance. 1,590 subjects (915 cases, 675 controls) in the general research use (GRU) group and 1,133 subjects (431 cases and 702 controls) in the autoimmune disease only (ADO) group were analyzed.

For the GRU group and ADO group, SNPs that failed to pass the HWE exact test were filtered (we used the p=0.001 as the threshold). Fisher exact test was used to compute the p-value of allele frequency for each SNP. The threshold of p-value was set as 1.2×10^−7^ by Bonferroni Correction. Then we selected first 100 SNPs of lowest p-values for further analysis. Two different large public datasets, gnomad. genome. NFE (Non-Finnish European, N = 7,509) (ref1) (19) and gnomad.exome.NFE (Non-Finnish European, N = 55,860) (ref2) (19), were selected as the reference groups. So there were four combinations: GRU group vs ref1 (GRU_ref1), GRU group vs ref2 (GRU_ref2), ADO group vs ref1 (ADO_ref1) and ADO group vs ref2 (ADO_ref2). For each condition, we computed the p-value of our model (see Additional files 3). Specially for the exome dataset (ref2), some SNPs not in the exome were excluded from the table.

To inspect the improvement of p-values in the whole level, we drew the Manhattan plot of GRU group (Figure 2) and ADO group (Figure 3) respectively. We observed notable changes of p-values before and after performing our method. The peaks of Figure 2 and Figure 3 jumped from about 1e-22 and 1e-32 to 1e-62 and 1e-52 respectively. Also the amount of SNPs with p-value between 1e-5 and 1e-8 has increased. The positive SNPs (*rs12191877*, *rs9468933*, etc.) became more positive due to the added genetic information of reference. Also, our method rescued a few SNPs which turned from negative to positive (see Additional files 4). Then, we searched Pubmed literature and found some SNPs associated with psoriasis that had been reported by other studies. By integrating the results of ref1 and ref2, we found that the SNPs with ref2 as reference were included in the results of ref1. And we presented the novel SNPs of GRU group (Table 2) and ADO group (Table 3). We calculated the genomic inflation factor, also known as lambda (λ). In GRU group, λ of P(T) and *P* (*ca–co*) were 1.08 and 0.98 respectively. And in ADO group, λ of P(T) and *p*(*ca*–*co*)were 0.81 and 0.96, respectively.

**Figure 2.**
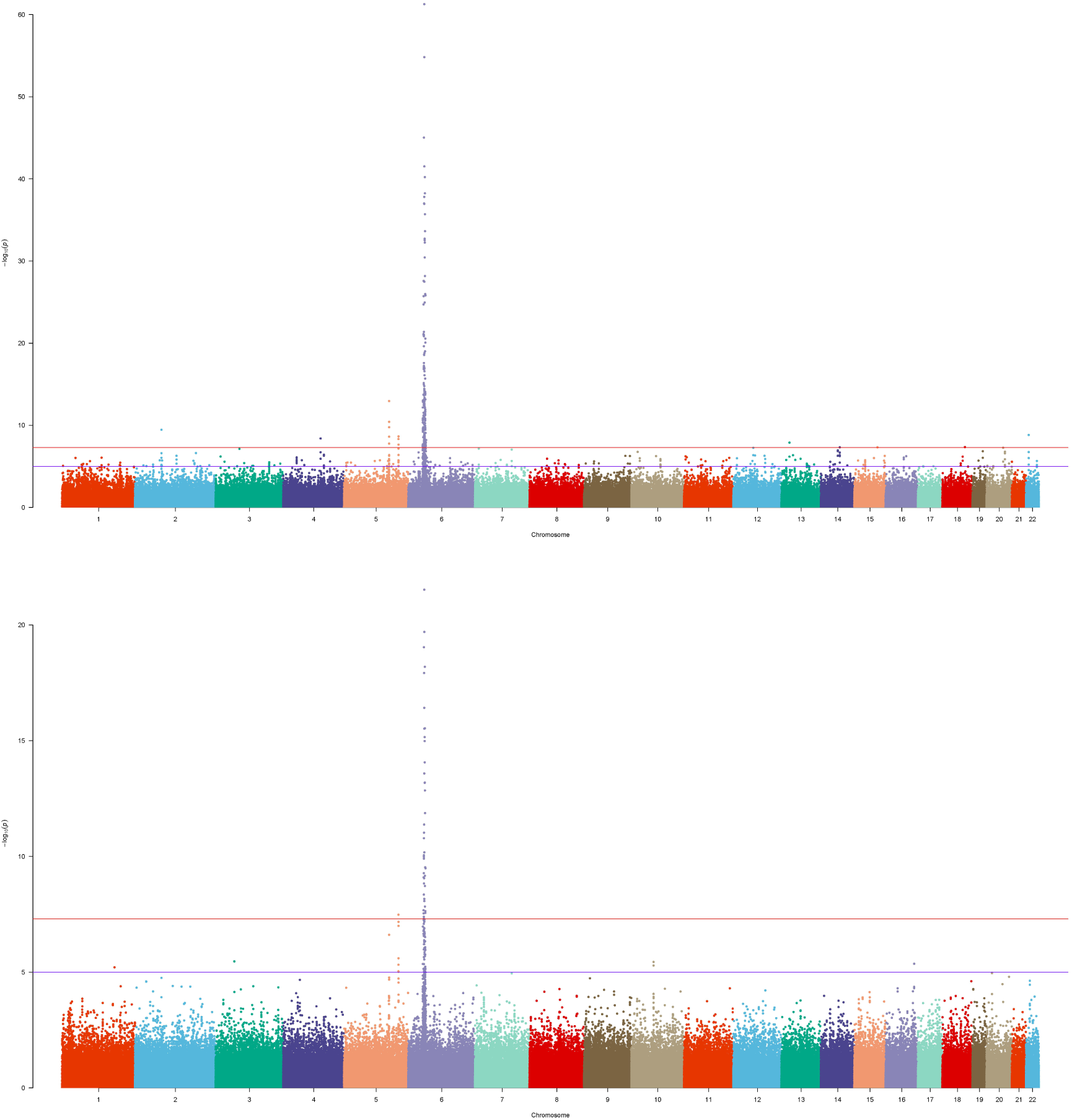
Manhattan plot of GRU group of Psoriasis GWAS datasets. The bottom figure corresponded to the P(ca-co) and top figure corresponded to the P(T), and reference was ref1.

**Table 2.** SNPs rescued from negative to positive of GRU group.

| RS | P(co- | P(ca-co) | P(T) | RS | P(co- | P(ca-co) | P(T) |
| --- | --- | --- | --- | --- | --- | --- | --- |
| *rs2021723* | 0.0943 | 1.66E-07 | 7.99E-22 | *rs9266825* | 0.79669 | 2.03E-07 | 1.25E-13 |
| *rs1015465* | 0.0777 | 2.22E-07 | 9.45E-22 | *rs9266845* | 0.74748 | 2.42E-07 | 2.50E-13 |
| *rs6906662* | 0.2178 | 3.06E-07 | 9.88E-20 | *rs9295991* | 0.74785 | 2.54E-07 | 3.28E-13 |
| *rs3094205* | 0.1810 | 1.76E-07 | 2.35E-19 | *rs9266813* | 0.97496 | 1.37E-06 | 3.87E-13 |
| *rs9262498* | 0.0562 | 2.28E-06 | 2.62E-18 | *rs9266846* | 0.69923 | 2.37E-07 | 4.00E-13 |
| *rs9262492* | 0.0782 | 2.24E-06 | 5.84E-18 | ***rs1051792*** | 0.94995 | 1.67E-06 | 4.77E-13 |
| *rs9295924* | 0.1989 | 1.04E-06 | 1.01E-17 | *rs176095* | 0.94513 | 2.04E-06 | 1.94E-12 |
| *rs6933779* | 0.2251 | 8.90E-07 | 1.15E-17 | *rs2239518* | 0.44369 | 1.48E-07 | 3.18E-12 |
| *rs2395471* | 0.4190 | 1.28E-07 | 1.37E-17 | *rs3130685* | 0.58732 | 5.00E-07 | 1.34E-11 |
| *rs10947208* | 0.4703 | 1.86E-07 | 2.65E-17 | *rs2894176* | 0.56223 | 6.25E-07 | 2.18E-11 |
| ***rs13437088*** | 0.5345 | 2.40E-07 | 8.09E-17 | ***rs2442719*** | 0.62654 | 1.07E-06 | 4.47E-11 |
| *rs4711229* | 0.6639 | 2.74E-07 | 7.27E-16 | *rs2734573* | 0.40816 | 4.93E-07 | 1.56E-10 |
| *rs2853950* | 0.8632 | 1.30E-07 | 1.70E-15 | *rs2858332* | 0.49528 | 1.11E-06 | 2.20E-10 |
| ***rs7192*** | 0.5438 | 8.41E-07 | 8.66E-15 | *rs3130048* | 0.15645 | 2.13E-07 | 2.73E-09 |
| *rs7194* | 0.5630 | 1.02E-06 | 1.25E-14 | *rs1003879* | 0.17704 | 4.68E-07 | 7.88E-09 |
| *rs12203586* | 0.2697 | 7.01E-06 | 1.27E-14 | *rs9391858* | 0.36543 | 6.00E-06 | 1.34E-08 |
| *rs2856726* | 0.3099 | 5.83E-06 | 1.90E-14 | *rs6894567* | 0.30040 | 4.80E-06 | 6.88E-08 |
| ***rs20541*** | 1 | 2.44E-07 | 1.13E-13 |  |  |  |  |

Note: The SNPs reported in pubmed were shown in bold.

**Table 3.**
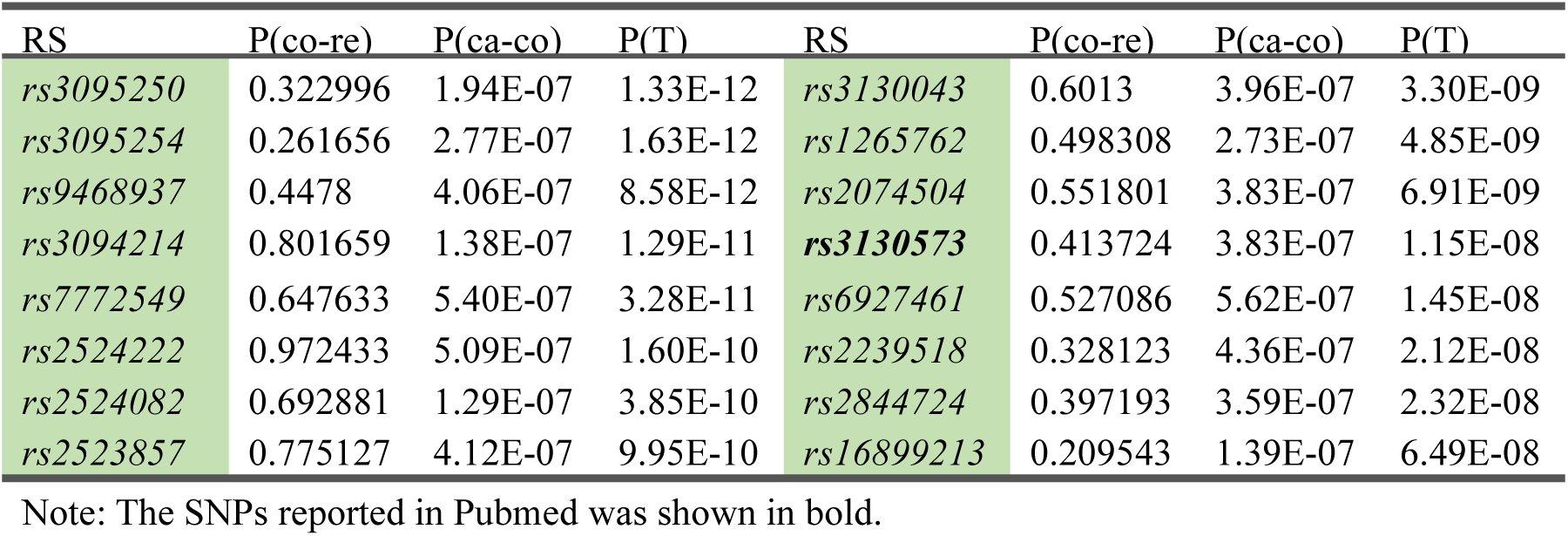
SNPs rescued from negative to positive of ADO group.

| RS | P(co-re) | P(ca-co) | P(T) | RS | P(co-re) | P(ca-co) | P(T) |
| --- | --- | --- | --- | --- | --- | --- | --- |
| *rs3095250* | 0.322996 | 1.94E-07 | 1.33E-12 | *rs3130043* | 0.6013 | 3.96E-07 | 3.30E-09 |
| *rs3095254* | 0.261656 | 2.77E-07 | 1.63E-12 | *rs1265762* | 0.498308 | 2.73E-07 | 4.85E-09 |
| *rs9468937* | 0.4478 | 4.06E-07 | 8.58E-12 | *rs2074504* | 0.551801 | 3.83E-07 | 6.91E-09 |
| *rs3094214* | 0.801659 | 1.38E-07 | 1.29E-11 | ***rs3130573*** | 0.413724 | 3.83E-07 | 1.15E-08 |
| *rs7772549* | 0.647633 | 5.40E-07 | 3.28E-11 | *rs6927461* | 0.527086 | 5.62E-07 | 1.45E-08 |
| *rs2524222* | 0.972433 | 5.09E-07 | 1.60E-10 | *rs2239518* | 0.328123 | 4.36E-07 | 2.12E-08 |
| *rs2524082* | 0.692881 | 1.29E-07 | 3.85E-10 | *rs2844724* | 0.397193 | 3.59E-07 | 2.32E-08 |
| *rs2523857* | 0.775127 | 4.12E-07 | 9.95E-10 | *rs16899213* | 0.209543 | 1.39E-07 | 6.49E-08 |

Note: The SNPs reported in Pubmed was shown in bold.

**Figure 3.**
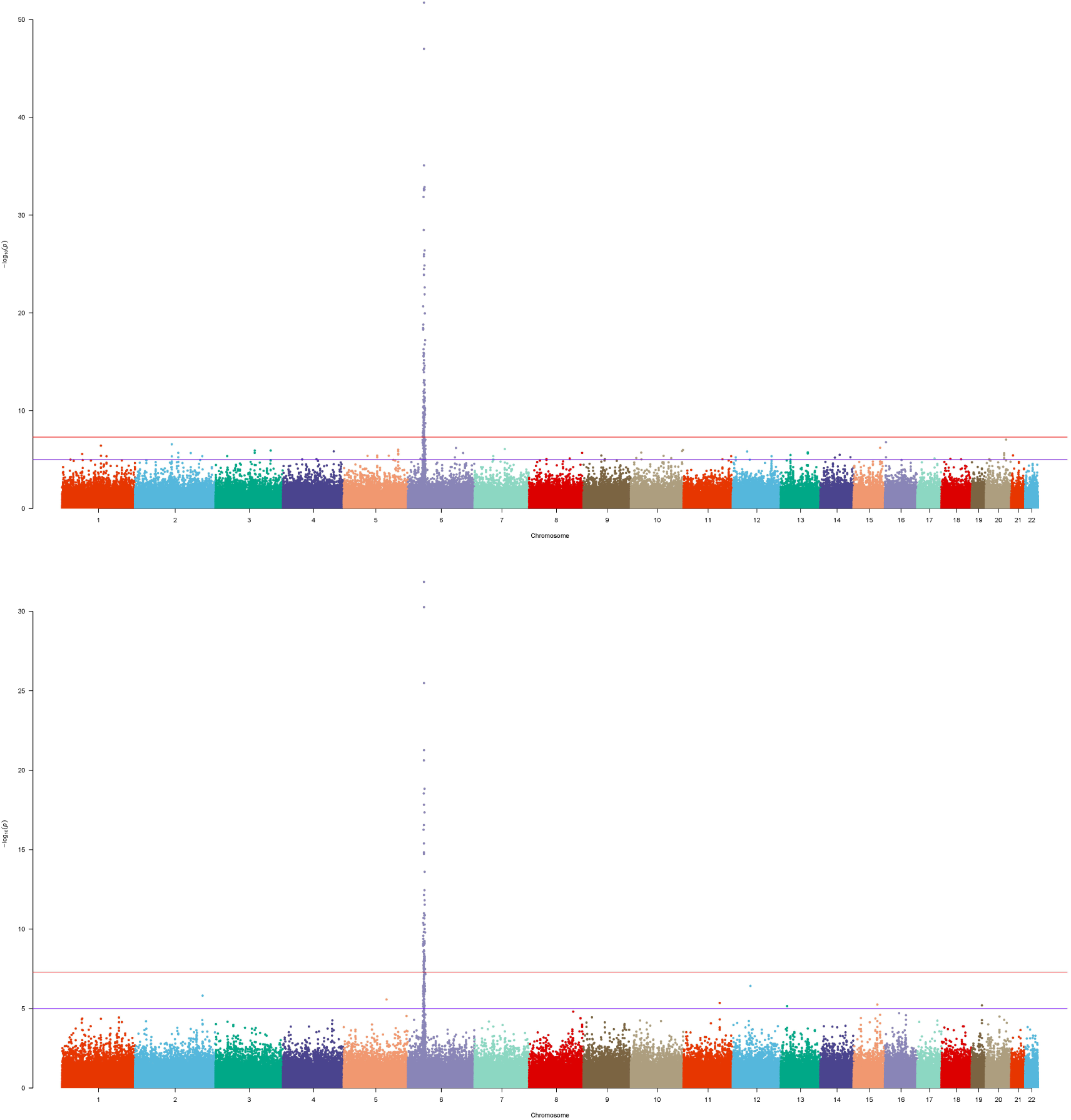
Manhattan plot of ADO group of Psoriasis GWAS datasets. The bottom figure corresponded to the P(ca-co) and top figure corresponded to the P(T), and reference was ref1.

For GRU group, *rs13437088* (P(T) = 8.09E-17), located 30 kb centromeric of *HLA-B* and 16 kb telomeric of *MICA* (MIM: 600169), had been previously reported to be associated with psoriasis (20). Besides, *rs7192* (P(T) = 8.66E-15) and *rs20541* (P(T) = 1.13E-13) were candidate causal SNPs at leukocyte antigen (HLA) loci (MIM: 142395) which played an important roles in pathways of psoriasis (21). Also, *rs1051792* (P(T) = 4.77E-13) in the *MICA* gene (rs1051792) had also been suggested to be specific for purely cutaneous manifestations of psoriasis (22). And SNP *rs2442719* (P(T) = 4.47E-11), located only 1kb from the telomeric end of *HLA-B* (MIM: 142830), also exhibited significant association with psoriasis (20). For the ADO group, *rs3130573* (P(T) = 1.70E-10) was located in *PSORS1C1* (MIM: 613525) gene which was a major susceptibility locus for psoriasis (23).

### Results from the schizophrenia GWAS datasets

We also applied our newly developed method to one schizophrenia GWAS (18) dataset to evaluate the performance. The SNPs that did not pass the HWE exact test were filtered (p=0.001). Fisher exact test was used to compute the p-value of allele frequency for each SNP. There was no statistically significant SNP with the threshold as 7.14×10^−8^ by Bonferroni Correction. The large public datasets, gnomad.genome.NFE (Non-Finnish European, N = 7509) (ref1) (19) was selected as reference to compute the p-value of our model (see Additional files 5).

For the schizophrenia GWAS dataset (18), the typical Fisher exact test did not yield genome-wide significant findings. After the introduction of reference, we found that several novel SNPs were associated with schizophrenia (Table 4). The Manhattan plot (Figure 4) clearly showed that before performing our method there were only a few SNPs with p-value below 1e-5 in the bottom figure. And after performing our method, plenty of SNPs came to the fore with p-value between 1e-5 and 1e-8. In addition, there were significant SNPs with p-value below 7e-8, some of which had been reported in previous studies. The SNP *rs12140791* is located in *NOS1AP* (MIM: 605551) gene which is essential for brain development and function and of potential relevance to schizophrenia (24). The *rs17021364* and *rs11097407* were reported to be associated with schizophrenia in a genome-wide meta-analysis (25). The *rs35648* (p-value = 9.65E-5) was also reported by a previous large-scale GWAS (26). The genomic inflation factors were 0.77 and 1.01 for P(T) and P(ca-co) respectively.

**Table 4.** SNPs rescued from negative to positive of schizophrenia dataset.

| RS | P(co-re) | P(ca-co) | P(T) | RS | P(co-re) | P(ca-co) | P(T) |
| --- | --- | --- | --- | --- | --- | --- | --- |
| ***rs12140791*** | 0.4979 | 1.92E-05 | 5.73E-10 | ***rs17021364*** | 0.62405 | 5.04E-05 | 1.64E-08 |
| *rs10753758* | 0.3999 | 3.84E-05 | 8.91E-10 | ***rs35648*** | 0.67746 | 5.29E-05 | 4.46E-08 |
| ***rs1109740777*** | 0.8144 | 2.03E-05 | 9.06E-09 |  |  |  |  |

Note: The SNPs reported in pubmed was shown in bold and italics.

**Figure 4.**
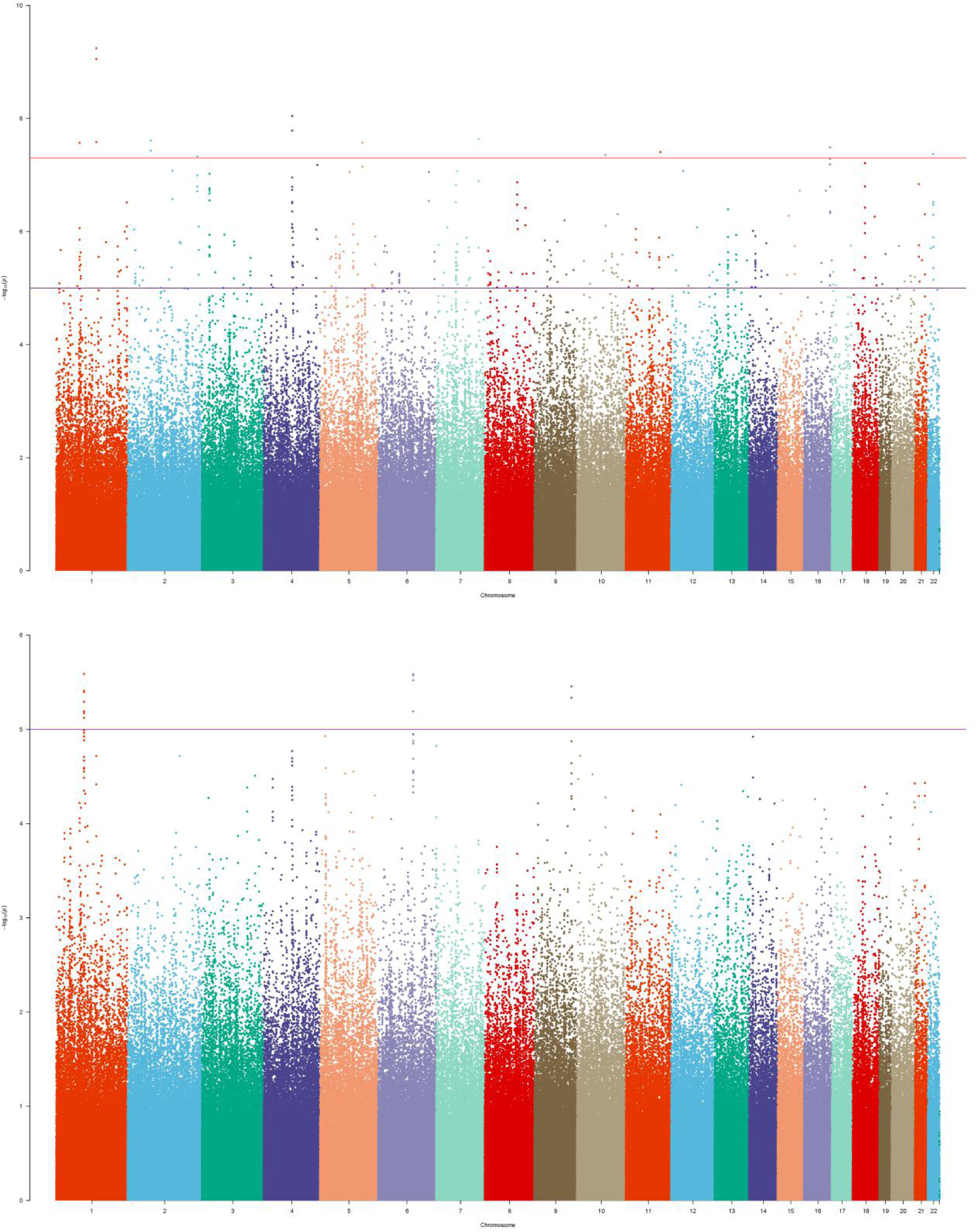
Manhattan plot of schizophrenia GWAS dataset. The bottom figure corresponded to the P(ca-co) and top figure corresponded to the P(T), and reference was ref1.

## DISCUSSION

Associations between SNPs and complex traits were detected by comparing frequencies of alleles in case and control group in GWAS (1). Several significant SNPs have been identified in classic GWAS studies (27-30). However, other SNPs of low frequencies which contribute to the complex traits remain hidden in the false negative results (31-35). To identify more novel susceptibility loci, large-scale GWAS is a costly and time-consuming approach. It is necessary to design more sensitive and accurate statistical methods. Interestingly, the results indicated that our method could increase the power which may contribute to detecting more significant SNPs.

Our simulations focused on several representative cases to evaluate the performance of the new model. When the allele frequencies of case, control and reference were equal, the power of T was higher than power of ca-(co+re), while both less than false positive rate. False positive results would be produced in the presence of small population genetic differentiation between control and reference. However, as the control size increased, the false positive rate of P(T) would be reduced (Table 1). The false positive rate would be controlled well, when the control sample was large enough. What’s more, T’s power was always higher than power of ca-co, when the allele frequencies of case and control were different, indicating that our method was more robust and had high sensitivity for GWAS.

For the psoriasis genome-wide association (GWAS) datasets (16,17), the positive SNPs became more positive and some of the negative SNPs turned to be positive after application of our method. The rescued SNPs, *rs1343708* (20), *rs7192* (21), *rs20541* (21), *rs1051792* (22), *rs2442719* (20) and *rs3130573* (23) were identified to be true positive. For the schizophrenia GWAS dataset (18), typical Fisher Exact tests produced no significant positive genetic loci. However, our method found that SNPs *rs12140791* (24), *rs10753758*, *rs11097407* (25), *rs17021364* (25), *rs35648* (26) could potentially be associated with schizophrenia. The results indicated that our method could be sensitive to generate more positive SNPs.

In the aforementioned application of our method to the psoriasis genome-wide association (GWAS) genetic datasets (16,17), we selected two large public datasets as reference groups (19). With different references, the model will compute different p-values for the test statistic. In the case of scenario, the p-value is lower than threshold of 0.01, indicating significant differentiation between control and reference, the result would be false positive. In this situation, selecting an appropriate dataset as the reference is the key to obtain better result. In addition, selecting multiple different references is feasible with the online tool. By comparing and integrating the output of different references, reasonable significant P(T) could provide more effective information for SNPs associated with the traits.

Novel SNPs with weak positive signals could be discovered when the sample size of case and control are insufficient in GWAS (2). With the support of reference database, the new p-value of some false negative genetic loci would decrease significantly down to the threshold. And our test statistic T is more robust than *p*(*ca*–(*co*+*re*)) as it takes the control-reference difference into account by penalization of *p*(*co*–*re*).

To conclude, the new statistic proposed here is effective to discover novel genome-wide significant loci with both small and large sample sizes.

## ACKNOWLEDGEMENT

We thank the Fudan University High-End Computing Center for supporting computations involved in this study.

## DATA AVAILABILITY

The psoriasis datasets and the schizophrenia dataset can be obtained from the Genetic Association Information Network (GAIN) database. The dbGaP accession number of the psoriasis datasets is phs000019.v1. p1 and the dbGaP access number of the schizophrenia dataset is phs000021.v1.p1. The online tool of Robust Reference Powered Association Test is available at http://drwang.top/gwas.html.

## SUPPLEMENTARY DATA

Supplementary Data are available at NAR online.

## REFERENCES

1. McCarthy MI, Abecasis GR, Cardon LR, Goldstein DB, Little J, Ioannidis JP, Hirschhorn JN. (2008). Genome-wide association studies for complex traits: consensus, uncertainty and challenges. NAT REV GENET., 9, 356–69.

2. He Y, Xu S, Jia C, Jin L. (2009). A design of multi-source samples as a shared control for association studies in genetically stratified populations. CELL RES., 19, 913–5.

3. Hakonarson H, Grant SF, Bradfield JP, Marchand L, Kim CE, Glessner JT, Grabs R, Casalunovo T, Taback SP, Frackelton EC, Lawson ML, et al. (2007). A genome-wide association study identifies KIAA0350 as a type 1 diabetes gene. NATURE, 448, 591–4.

4. Zeggini E, Weedon MN, Lindgren CM, Frayling TM, Elliott KS, Lango H, Timpson NJ, Perry JR, Rayner NW, Freathy RM, Barrett JC, et al. (2007). Replication of genome-wide association signals in UK samples reveals risk loci for type 2 diabetes. SCIENCE, 316, 1336–41.

5. Parkes M, Barrett JC, Prescott NJ, Tremelling M, Anderson CA, Fisher SA, Roberts RG, Nimmo ER, Cummings FR, Soars D,et al. (2007). Sequence variants in the autophagy gene IRGM and multiple other replicating loci contribute to Crohn’s disease susceptibility. NATURE, 39, 830–2.

6. Thomas G, Jacobs KB, Yeager M, Kraft P, Wacholder S, Orr N, Yu K, Chatterjee N, Welch R, Hutchinson A, et al. (2008). Multiple loci identified in a genome wide association study of prostate cancer. Nature Genet., 40, 310–5.

7. Easton DF, Pooley KA, Dunning AM, Pharoah PD, Thompson D, Ballinger DG, Struewing JP, Morrison J, Field H, Luben R, et al. (2007). Genome-wide association study identifies novel breast cancer susceptibility loci. NATURE, 447, 1087–93.

8. Wellcome Trust Case Control Consortium. (2007). Genome-wide association study of 14,000 cases of seven common diseases and 3,000 shared controls. NATURE, 447, 661–78.

9. Bacanu SA, Devlin B, Roeder K. (2000). The power of genomic control. AM J HUM GENET., 66, 1933–44.

10. He Y, Jiang R, Fu W, Bergen AW, Swan GE, Jin L. (2008). Correlation of population parameters leading to power differences in association studies with population stratification. ANN HUM GENET., 72, 801–11.

11. R. A. Fisher. (1922). On the interpretation of χ2 from contingency tables, and the calculation of P. Journal of the Royal Statistical Society, 85, 87–94.

12. R. A. Fisher. (1954). Statistical Methods for Research Workers. Oliver and Boyd.

13. Agresti A. (1992). A Survey of Exact Inference for Contingency Tables. STAT SCI., 7, 131–53.

14. Casella G, Berger RL. (2002). Statistical Inference: Thomson Learning.

15. Mcneil AJ. (2003). The Laplace Distribution and Generalizations: A Revisit With Applications to Communications, Economics, Engineering, and Finance. Journal of the Royal Statistical Society, 52, 698–9.

16. Fang S, Fang X, Xiong M.(2011). Psoriasis prediction from genome-wide SNP profiles. BMC Dermatol., 11, 1.

17. Nair RP, Stuart PE, Nistor I, Hiremagalore R, Chia NV, Jenisch S, Weichenthal M, Abecasis GR, Lim HW, Christophers E, Voorhees JJ, Elder JT, et al. (2006). Sequence and haplotype analysis supports HLA-C as the psoriasis susceptibility 1 gene. AM J HUM GENET., 75, 827–51.

18. Zuo L, Wang K, Zhang XY, Pan X, Wang G, Tan Y, Zhong C, Krystal JH, State M, Zhang H, Luo X. (2013). Association between common alcohol dehydrogenase gene (ADH) variants and schizophrenia and autism. HUM GENET., 132, 735–43.

19. Lek M, Karczewski KJ, Minikel EV, Samocha KE, Banks E, Fennell T, O’Donnell-Luria AH, Ware JS, Hill AJ, Cummings BB, et al. (2016). Analysis of protein-coding genetic variation in 60,706 humans. NATURE, 536, 285–91.

20. Feng BJ, Sun LD, Soltaniarabshahi R, Bowcock AM, Nair RP. (2009). Multiple loci within the major histocompatibility complex confer risk of psoriasis. PLOS GENET., 5, e1000606.

21. Lee YH, Choi SJ, Ji JD, Song GG. (2012). Genome-wide pathway analysis of a genome-wide association study on psoriasis and Behcet’s disease. MOL BIOL REP., 39, 5953–9.

22. Bowes J, Budu-Aggrey A, Huffmeier U, Uebe S, Steel K, Hebert HL, Wallace C, Massey J, Bruce IN, Bluett J, et al. (2015). Dense genotyping of immune-related susceptibility loci reveals new insights into the genetics of psoriatic arthritis. NAT COMMUN., 6, 6046.

23. Fan X, Yang S, Huang W, Wang ZM, Sun LD, Liang YH, Gao M, Ren YQ, Zhang KY, Du WH, Shen YJ, Liu JJ, Zhang XJ. (2008). Fine mapping of the psoriasis susceptibility locus PSORS1 supports HLA-C as the susceptibility gene in the Han Chinese population. PLOS GENET., 4, e1000038.

24. Glessner JT, Reilly MP, Kim CE, Takahashi N, Albano A, Hou C, Bradfield JP, Zhang H, Sleiman PM, Flory JH, et al. (2010). Strong synaptic transmission impact by copy number variations in schizophrenia. Proc Natl Acad Sci U S A., 107, 10584–9.

25. Wang KS, Liu XF, Aragam N. (2010). A genome-wide meta-analysis identifies novel loci associated with schizophrenia and bipolar disorder. SCHIZOPHR RES., 124, 192–9.

26. Shi J, Levinson DF, Duan J, Sanders AR, Zheng Y, Pe’er I, Dudbridge F, Holmans PA, Whittemore AS, Mowry BJ, et al. (2009). Common variants on chromosome 6p22.1 are associated with schizophrenia. NATURE, 460, 753–7.

27. Scott LJ, Mohlke KL, Bonnycastle LL, Willer CJ, Li Y, Duren WL, Erdos MR, Stringham HM, Chines PS, Jackson AU, et al. (2007). A genome-wide association study of type 2 diabetes in Finns detects multiple susceptibility variants. SCIENCE, 316, 1341–5.

28. Duerr RH, Taylor KD, Brant SR, Rioux JD, Silverberg MS, Daly MJ, Steinhart AH, Abraham C, Regueiro M, Griffiths A, et al. (2006). A genome-wide association study identifies IL23R as an inflammatory bowel disease gene. SCIENCE, 314, 1461–3.

29. Hunter DJEA. (2007). A genome-wide association study identifies alleles in FGFR2 associated with risk of sporadic postmenopausal breast cancer. Nature Genet., 39, 870–4.

30. Hunter DJ, Kraft P, Jacobs KB, Cox DG, Yeager M, Hankinson SE, Wacholder S, Wang Z, Welch R, Hutchinson A, et al. (2007). A common allele on chromosome 9 associated with coronary heart disease. SCIENCE, 316, 1488–91.

31. Willer CJ, Sanna S, Jackson AU, Scuteri A, Bonnycastle LL, Clarke R, Heath SC, Timpson NJ, Najjar SS, Stringham HM, et al. (2008). Newly identified loci that influence lipid concentrations and risk of coronary artery disease. Nature Genet., 40, 161–9.

32. Sanna S, Jackson AU, Nagaraja R, Willer CJ, Chen WM, Bonnycastle LL, Shen H, Timpson N, Lettre G, Usala G, et al. (2008). Common variants in the GDF5-UQCC region are associated with variation in human height. Nature Genet., 40, 198–203.

33. Frayling TM, Timpson NJ, Weedon MN, Zeggini E, Freathy RM, Lindgren CM, Perry JR, Elliott KS, Lango H, Rayner NW, et al. (2007). A common variant in the FTO gene is associated with body mass index and predisposes to childhood and adult obesity. SCIENCE, 316, 889–94.

34. Nair RP, Duffin KC, Helms C, Ding J, Stuart PE, Goldgar D, Gudjonsson JE, Li Y, Tejasvi T, Feng BJ, et al. (2009). Genome-wide scan reveals association of psoriasis with IL-23 and NF-κB pathways. NAT GENET., 41, 199–204.

35. Wang WJ, Yin XY, Zuo XB, Cheng H, Du WD, Zhang FY, Yang S, Zhang XJ. (2013). Gene– gene interactions in IL23/Th17 pathway contribute to psoriasis susceptibility in Chinese Han population. J EUR ACAD DERMATOL., 27, 1156–62.

